# Tissue-specific I-Smad mechanisms revealed by structure-function analysis in *Drosophila*

**DOI:** 10.1101/2025.06.20.660739

**Authors:** Ania M. Simoncek, Steven J. Sviridoff, Joshua N. Hays, Noah J. Graichen, Mikolaj J. Sulkowski

## Abstract

Inhibitory Smads (I-Smads) regulate TGF-β/BMP signaling through multiple distinct mechanisms, but whether different tissues preferentially employ specific mechanisms remains unknown. To address this question, we performed structure-function analyses of the *Drosophila* I-Smad Dad and its vertebrate orthologs Smad6 and Smad7 in neural and wing tissues, measuring in vivo outputs of BMP signaling. We identified a critical 24-amino acid putative DNA-binding domain (DNABD) within the MH1 domain of the *Drosophila* I-Smad, Dad, that is essential for inhibitory function in wing tissue but unessential in neural tissue. Structural analyses revealed that ΔDNABD disrupts a beta hairpin structure homologous to R-Smad DNA-binding regions. We also found that Dad requires an intact MH1 domain to disrupt wing development, whereas either MH1 or MH2 can independently disrupt BMP signaling in motor neurons. These findings support a model where Dad functions through MH1-mediated transcriptional regulation in wing primordium, but through multiple mechanisms in neurons. Comparative analysis revealed that vertebrate I-Smad orthologs also show tissue-specific activity patterns, with structural predictions suggesting that Smad6 retains ancestral DNA-binding capacity while Smad7 has evolved enhanced MH2-mediated functions. These results reveal context-dependent mechanisms of I-Smads that further the understanding of TGF-β/BMP pathway regulation.

## Introduction

Transforming growth factor β (TGF-β) signaling is an ancient and evolutionarily conserved pathway for intercellular communication that controls diverse life processes such as axis formation, cell differentiation, and tissue homeostasis. Its pleiotropy necessitates precise intra- and extracellular modulation to elicit the appropriate effects in target cells (Massagué 2012). Dysregulation of TGF-β signaling underlies the pathophysiology of numerous disease states and developmental disorders. For instance, as TGF-β inhibits mitosis and promotes apoptosis in many tissues, loss of TGF-β signaling often engenders tumorigenesis. In later oncogenic stages, however, cells become resistant to the apoptosis-inducing effects, and TGF-β promotes tumor growth and invasiveness via dedifferentiation, angiogenesis, and immune suppression (Derynck et al. 2001; Syed 2016; Chan et al. 2022). Thus, the ability of cells to receive and interpret physiologically appropriate levels of TGF-β signaling is vital across metazoan life stages.

Inhibitory Smads (I-Smads) are intracellular proteins that ensure an appropriate response to TGF-β signaling. They are transcriptional targets of the signaling pathway, functioning in a negative-feedback mechanism (Miyazawa and Miyazono 2017). Dysfunction of I-Smads is implicated in the pathogenesis of human diseases such as colorectal cancer, atrial fibrosis, and inflammatory bowel disease (Broderick et al. 2007; Fortini et al. 2014; He et al. 2011; Tang et al. 2019). Although mammalian genomes encode two I-Smads (*Smad6* and *Smad7*), the *Drosophila* genome contains only a single I-Smad encoding gene, *Daughters against Decapentaplegic* (*Dad*) (Tsuneizumi et al. 1997; Massagué et al. 2005). This lack of genetic redundancy, combined with powerful genetic tools and well-characterized phenotypes, make *Drosophila* an ideal system in which to study the mechanisms of I-Smad function *in vivo*.

I-Smads belong to the Smad family of proteins, which transduce signals from TGF-β receptors. The Smad family also includes R-Smads, which are the receptor-regulated Smads that become activated by type I receptors, and Co-Smads (common-mediator Smads), which form complexes with activated R-Smads to enable their nuclear translocation and transcriptional activity (Schmierer and Hill 2007). Vertebrate R-Smads include Smad2 and Smad3, which respond to TGF-β and activin signals, and Smad1, Smad5, and Smad8/9, which respond to BMP signals (Massagué et al. 2005). Smads are characterized by the presence of two main functional domains: the MH1 domain, which mediates DNA binding and nuclear import, and the MH2 domain, which facilitates protein-protein interactions, including receptor binding, Smad oligomerization, and transcriptional activation. These two domains are connected by a variable linker region that contains regulatory sites for post-translational modifications.

I-Smads share a similar domain organization with other Smads. The crystal structure of SMAD7 reveals that the MH2 domain shares a high degree of structural similarity with the MH2 domains of other Smads (Murayama et al. 2020). However, there is currently no crystal structure available for the I-Smad N-terminal region, and no consensus on whether it constitutes a true MH1 domain. The two mammalian I-Smad paralogs, Smad6 and Smad7, are divergent both in their structure and function. Smad7 inhibits both TGF-β and bone morphogenetic protein (BMP) signaling, whereas Smad6 specifically inhibits BMP signaling (Hanyu et al. 2001; Goto et al. 2007). Smad7 is also more potent than Smad6 when repression is quantified via luciferase assay (Hanyu et al. 2001). Previous studies, largely performed in vitro, indicate that Dad specifically inhibits the BMP branch of TGF-β signaling, and this inhibition has been attributed to Dad preventing Thickveins, a BMP type I receptor, from phosphorylating the R-Smad Mothers against decapentaplegic (Mad) (Inoue et al. 1998; Kamiya et al. 2008; Li et al. 2017).

Intriguingly, I-Smads have been shown to antagonize TGF-β signaling by several distinct mechanisms in addition to the abovementioned inhibition of type I receptors (Miyazawa and Miyazono 2017). I-Smads can form complexes with R-Smads, thus preventing their association with co-Smads (Hata and Chen 2016; Yan et al. 2016). I-Smads cooperate with E3 ubiquitin ligases to target R-Smads and other TGF-β components for destruction (Zhu et al. 1999; Kavsak et al. 2000; Ebisawa et al. 2001; Murakami et al. 2003; Morén et al. 2005). I-Smads can also accumulate in the nucleus and directly regulate transcription to counter the effects of R-Smads (Bai et al. 2000; Hanyu et al. 2001; Zhang et al. 2007). Given that I-Smads function through such diverse mechanisms and considering that many of the previous studies were performed in vitro, we sought to further explore the mechanisms of Dad and I-Smad orthologs by conducting structure/function studies in different in vivo cellular contexts.

Using a series of transgenic animals, here we provide evidence that the human orthologs of Dad, SMAD6 and SMAD7, have diverging functions in *Drosophila* wing and neural tissue. We further demonstrate that Dad lacking a specific, putative DNA binding region of its MH1 domain is not sufficient to disrupt wing development yet is sufficient to inhibit BMP signaling in neural tissue. By comparing protein structure predictions, we extend these results to hypothesize that Smad6/Dad inhibit TGF-β signaling by a divergent mechanism from Smad7, and that this mechanism necessitates the MH1 domain, possibly for DNA binding. We also find that in both brain and wing tissue, Dad does not require palmitoylation at a C-terminal cysteine (C556) previously shown to be required for inhibition in ovarian tissue (Li et al. 2017). Together these findings shed light on the molecular mechanisms of TGF-β regulation and provide compelling evidence for the evolution of distinct, tissue-specific functionality among I-Smad orthologs.

## Results

### Creation of a series of transgenic animals to investigate tissue-specific I-Smad functionality

*T*o investigate the structural basis for alternative I-Smad mechanisms, we created a series of transgenic *Drosophila* to express vertebrate I-Smad orthologs, Smad6 and Smad7, and targeted alterations of the *Dad* gene (Figure 1A). In addition to full-length *Smad6* and *Smad7*, a transgenic line expressing a shorter isoform of *Smad7* lacking the MH1 domain (*Smad7S*) was developed. Dad truncations covering the MH1 and MH2 domains were constructed, along with truncations deleting specific residues implicated in DNA binding and trimerization (Δ*DNABD* and *ΔTrimer*). We also created a substitution replacing a C-terminal cysteine of Dad that is a reported target of palmitoylation (Dad^C556A^).

**Figure 1:**
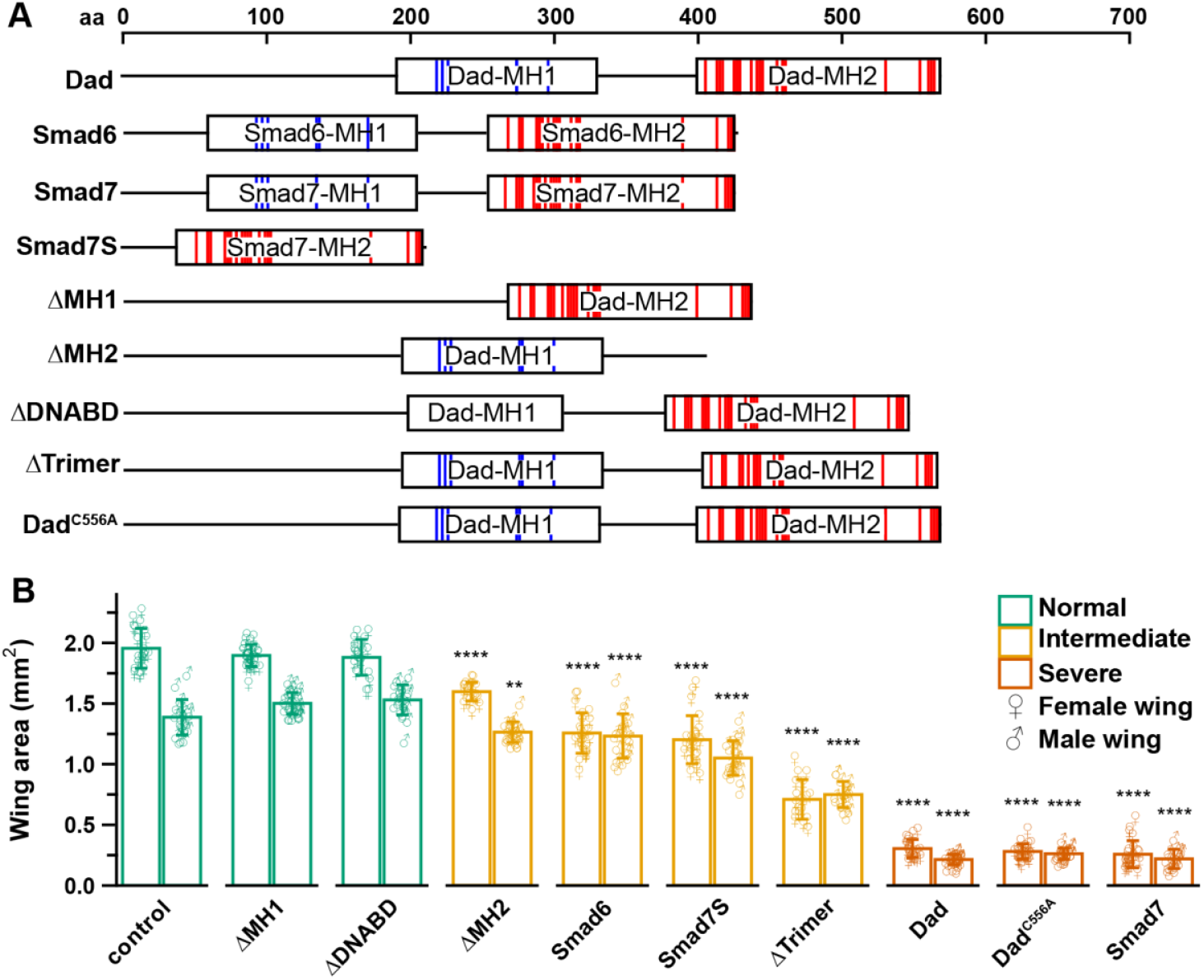
Disruption of *Drosophila* wing development by I-Smad structure/function constructs and vertebrate orthologs. (A) Protein domain organization of I-Smad constructs. Scale indicates length in amino acids. Locations of putative DNA binding sites (blue) and putative trimer interface (red) are indicated. (B) Quantification of wing area (mm^2^) in adult flies expressing I-Smad constructs in wing primordium with *Ap-Gal4* and genetic background controls (*Ap-Gal4/+*). Data points for individual female or male wings are represented by ♀ or ♂ symbols, respectively. Bars represent mean ± SD. Statistical comparisons performed using Dunnett’s test comparing individual genotypes with controls. **** p<0.0001, ** p<0.01

### Phenotypic analysis of wing tissue following construct expression

To assess the potency of our gene products to inhibit BMP signaling in developing wing tissue, constructs were expressed with *Ap-Gal4*, which is expressed primarily in the dorsal compartment of the wing imaginal disc (O’Keefe et al., 1998, Butterworth & King, 1965). We found that phenotypes clustered into three groups based on severity. Group 1, which includes genetic background control, ΔMH1, and ΔDNABD, had wing phenotypes indistinguishable from controls. The mean wing areas (in mm^2^) for males were 1.39±0.146, 1.50±0.089 and 1.53±0.124, and for females were 1.96±0.163, 1.90±0.091, and 1.88±0.147, respectively. Group 2 includes ΔMH2, Smad7S, Smad6, and ΔTrimer. This group had a moderately severe phenotypes, with mean areas for males of 1.26±0.085, 1.05±0.143, 1.23±0.183, and 0.749±0.107. Group 2 females wings were 1.60± 0.076, 1.20±0.198, 1.26±0.167, and 0.710±0.163, respectively. Finally, group 3, which includes Dad, Smad7L, and Dad^C556A^, had the most severe phenotypes with mean areas for males of 0.214±0.045, 0.219±0.079, and 0.262±0.047, and for females of 0.304±0.076, 0.258±0.112, and 0.280±0.063, respectively (Figure 1B).

Based upon these results, we conclude that Dad lacking its MH1 domain is insufficient to inhibit wing tissue growth. Dad with a 24 aa deletion in a putative DNA binding site (ΔDNA-BD) within the MH1 domain is also insufficient. Conversely, Smad7 lacking the MH1 domain is sufficient to inhibit wing growth. Even with a substitution converting the cysteine at position 556 to alanine (C556A), Dad remained effective in inhibiting wing growth. These results indicate that Dad must include MH1 domain to disrupt wing growth, while the vertebrate counterpart Smad7 can disrupt growth without the MH1 domain. Furthermore, Dad lacking the MH2 domain is still capable of producing some disruption in wing development.

### Structural analysis of MH1 domains

As the MH1 domain of Dad was sufficient to disrupt wing development, we sought to identify potential differences in protein structures among the MH1 domains of I-Smad orthologs. We first generated a model of the MH1 domain of the R-Smad, Smad3 using AlphaFold 3 (Figure 2A). The model matched the previously published crystal structure (Shi et al. 1998). Notably, the β-hairpin structure and its interaction with DNA were correctly replicated. When we compared predictions of the MH1 structures of Dad, Smad6, and Smad7 we found that the β-hairpin structure is longer and more disorganized in Smad7 compared with Dad or Smad6, which had β-hairpin structures with antiparallel stretches that more closely resemble the DNA binding structure in Smad3 (Figure 2B).

**Figure 2:**
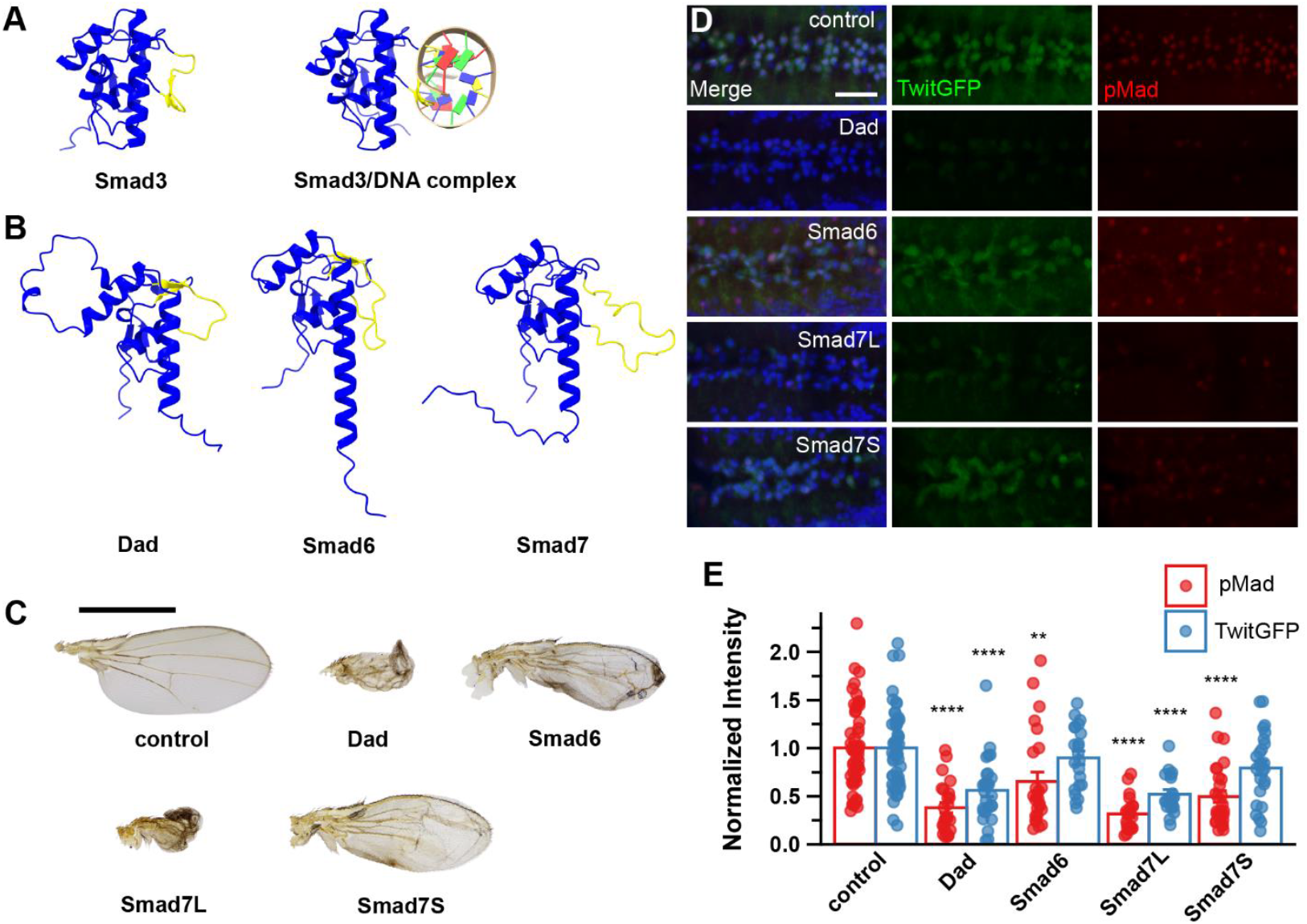
Human I-Smads inhibit BMP signaling in *Drosophila* tissues. (A-B) AlphaFold3 structural predictions of MH1 domains. Beta hairpin region shown in yellow. Remaining MH1 shown in blue. (A) Smad3 MH1 domain showing DNA binding region and predicted interaction. (B) Dad, Smad6, and Smad7 MH1 domains. (C) Representative wings from animals expressing indicated construct in wing primordium. Scale bar represents 1 mm. (D-E) Transgenic control (*TwitGFP, OK371-Gal4/+*) compared with *OK371-Gal4*-driven overexpression of *UAS-Dad, UAS-Smad6, UAS-Smad7L*, and *UAS-Smad7S*. (D) Motor neurons in ventral nerve cord triple-labeled for Elav (blue), Twit-GFP (green) and pMad (red). Scale bar represents 20 microns. (E) Quantification of pMad and Twit-GFP intensity normalized to genetic background control. Bars represent mean ± SEM. Each dot represents mean data from motor neurons of one CNS. Statistical comparisons performed using Dunn’s test with Bonferroni correction. **** p<0.0001, ** p<0.01.

**Figure 3:**
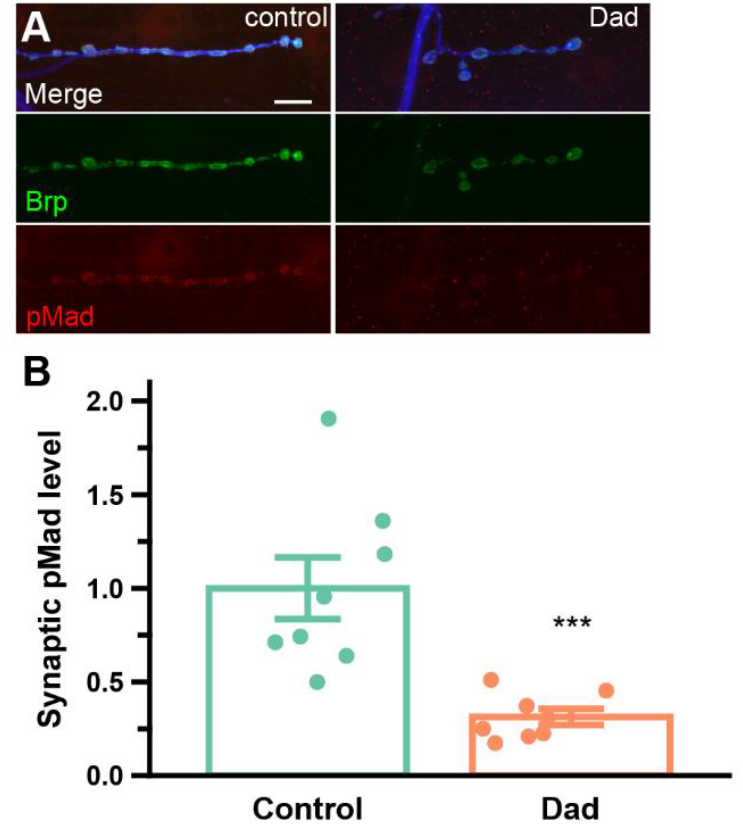
Presynaptic Dad expression inhibits local BMP signaling at NMJ synapses. Transgenic control (*TwitGFP, OK371-Gal4/+*) compared with *OK371-Gal4*-driven overexpression of *UAS-Dad*. (A) Muscle 4 NMJ triple-labeled for the presynaptic membrane marker HRP (blue), active zone marker Brp (green), and synaptic pMad (red). Scale bar represents 10 microns. (B) Quantification of synaptic pMad intensity. Each dot represents one data point, which is the mean of one animal’s NMJ 4s from A2 and A3 normalized to control (4 NMJs per data point). Bars represent mean ± SD. Statistical comparison performed with Student’s t test. *** p<0.005.

### Effects of human I-Smad orthologs in neural tissue

To gauge the ability of specific I-Smad structural regions to inhibit BMP signaling in motor neurons, we expressed constructs using *OK371-Gal4*, a Gal4 enhancer trap in the proximity of the *DVGLUT* gene, which drives expression specifically in glutamatergic motor neurons (Mahr and Aberle 2006; Sanyal 2009). We began by determining the ability of the vertebrate I-Smads, Smad6 and Smad7, to inhibit BMP signaling in this neural tissue. BMP signaling was quantified by visualizing levels of phosphorylated Mad (pMad) in motor nuclei. Smad6, Smad7L, and Smad7S decreased nuclear pMad relative to genetic background control (mean±s.e.m.: 0.650±0.0.098, 0.315±0.040, and 0.495±0.061, respectively) (Figure 2). To evaluate a downstream effector of BMP signaling, we utilized a GFP protein trap in the BMP target gene *twit* (Sulkowski et al., 2016; Kim & Marquez, 2012; Venken et al., 2011). Smad7L decreased Twit-GFP levels to 0.519±0.0.048 relative to control. Interestingly, Smad6 and Smad7S did not cause a significant reduction in Twit-GFP levels (0.897±0.070 and 0.792±0.072 relative to control, respectively). Thus, vertebrate I-Smad orthologs were sufficient in inhibiting BMP signaling in neural as well as wing disc tissues.

### Regulation of non-canonical BMP signaling at neuromuscular synapses by I-Smads

BMP signaling at the neuromuscular junction (NMJ) encompasses both the canonical retrograde pathway, which culminates with pMad accumulation in the nucleus (Aberle et al. 2002; Marqués et al. 2002; McCabe et al. 2003), and a non-canonical pathway, characterized by pMad accumulation at synaptic active zones (Dudu et al. 2006; Higashi-Kovtun et al. 2010; O’Connor-Giles et al. 2008; Smith et al. 2012; Sulkowski et al. 2014, 2016). To test whether Dad affects the noncanonical BMP pathway, we examined synaptic pMad levels in animals overexpressing Dad in motor neurons. We saw a significant reduction in pMad accumulation at NMJ synapses in animals overexpressing Dad in motor neurons relative to controls (0.313±0.043; p<0.005). We also observed defects in NMJ morphology (bouton number and size) reminiscent of BMP pathway perturbations (data not shown). Therefore, I-Smads can inhibit both canonical and noncanonical BMP signaling at the NMJ.

### I-Smad MH1 and MH2 domain truncations in neural tissue

To evaluate the necessity of I-Smad structural domains to inhibit BMP signaling in motor neurons, expression of *Dad* constructs with truncations of the MH1 and MH2 domains (ΔMH1 and ΔMH2) were driven with *OK371-Gal4* (Figure 4). Both ΔMH1 and ΔMH2 were sufficient to suppress nuclear pMad accumulation (0.201±0.015 and 0.396±0.058 relative to control, respectively). ΔMH1 expression significantly reduced Twit-GFP reporter gene expression to 0.525±0.047 relative to control. ΔMH2 reduced Twit-GFP to 0.734±0.083, but this reduction was not statistically significant. This analysis reveals that the functional domains of *Dad* can independently inhibit BMP signaling in neural tissue, with the mechanisms utilizing the MH2 domain appearing to be slightly more effective in motor neurons. This contrasts with the results of expression in wing primordium tissue, where there was no difference observed when ΔMH1 was expressed and only a small reduction in area when ΔMH2 was expressed (Figure 4C).

**Figure 4:**
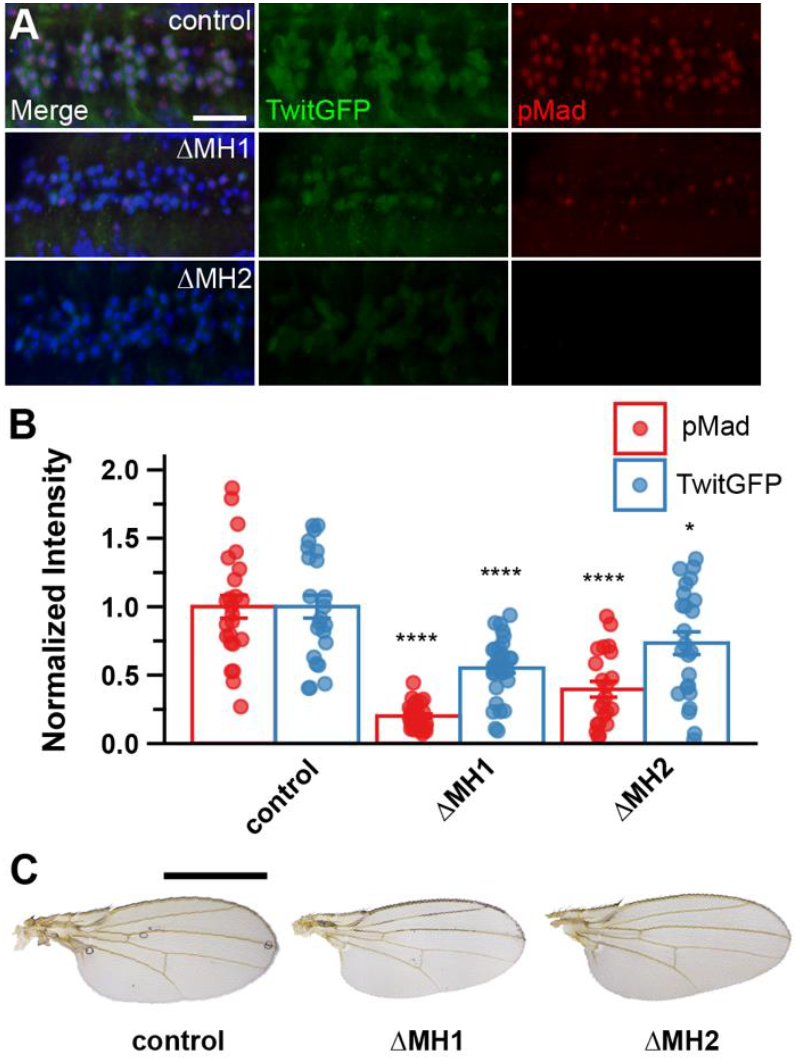
Dad can repress BMP signaling in Brains through multiple mechanisms. Transgenic control (*TwitGFP, OK371-Gal4/+*) compared with OK371-Gal4 driven overexpression of UAS-ΔMH1 and UAS-ΔMH2 (A) Motor neurons in ventral nerve cord triple-labeled for Elav (blue), Twit-GFP (green) and pMad (red). Scale bar represents 20 microns. (B) Quantification of pMad and Twit-GFP intensity. Bars represent mean ± SEM. (C) Representative images of wing phenotypes for indicated genotypes. Scale bar represents 1mm. Statistical comparisons performed using Dunn’s test with Bonferroni correction. **** p<0.0001, * p<0.05.

### Deletion of putative DNA binding sites and trimerization interfaces

To determine whether specific regions of the I-Smad MH1 and MH2 domains are sufficient to inhibit BMP signaling in motor neurons, we drove the expression of *Dad* constructs with deletions in putative DNA binding sites (ΔDNABD) or trimer interfaces (ΔTrimer) with OK371-Gal4 (Figure 5). Structural predictions show that the ΔDNABD mutation removes the beta-hairpin structure implicated in DNA binding (Figure 5A). Both ΔDNABD and ΔTrimer significantly reduced the accumulation of nuclear pMad relative to control (0.292±0.0276 and 0.341±0.0357, respectively). While ΔTrimer caused a significant decrease in Twit-GFP reporter gene levels (0.656±0.062), ΔDNABD expression did not significantly change Twit-GFP expression (0.849±0.0457). Therefore, Dad lacking putative DNA binding sites within the MH1 domain was not sufficient for repressing this target gene’s expression. However, this same structure is sufficient for decreasing pMad accumulation, indicating that these processes are separable and can be attributed to different structural motifs. As the ΔDNABD construct was not sufficient for altering wing morphology (Figures 1B and 5D), it appears that different tissue types preferentially employ different I-Smad mechanisms.

**Figure 5:**
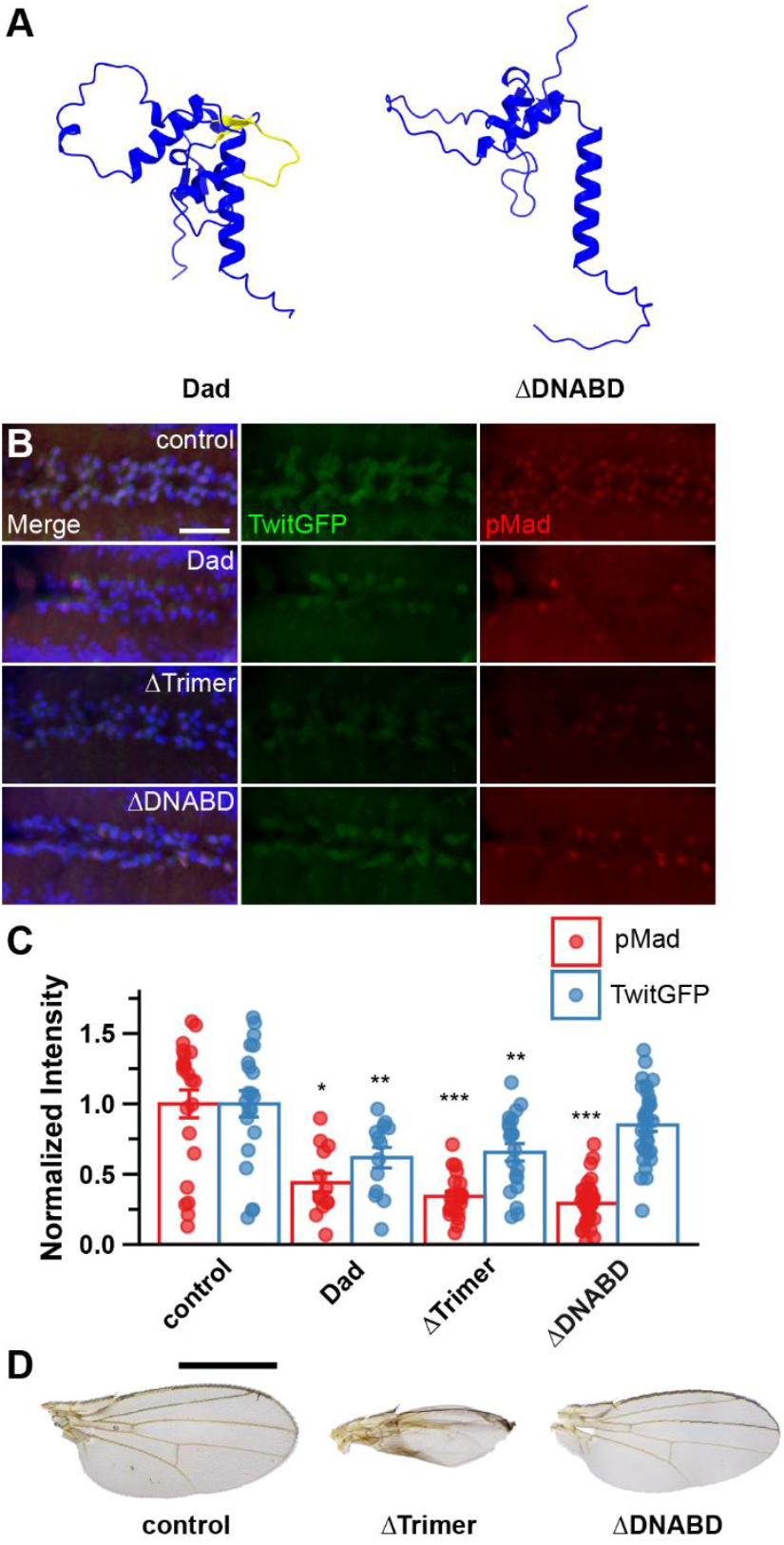
Dad lacking putative DNA binding region is sufficient for repressing pMad accumulation and Twit-GFP levels in brains, but not for repressing wing development. (A) AlphaFold3 structure prediction of Dad and Dad^ΔDNABD^ MH1 domains. Putative Dad DNA binding domain is depicted in yellow (absent in Dad^ΔDNABD^). (B-C) Transgenic control (*TwitGFP, OK371-Gal4/+*) compared with *OK371-Gal4*-driven overexpression of *UAS-Dad*^*ΔTrimer*^ and *UAS-Dad*^*ΔDNABD*^. (B) Motor neurons in ventral nerve cord triple-labeled for Elav (blue), Twit-GFP (green) and pMad (red). Scale bar represents 20 microns. (C) Quantification of pMad and Twit-GFP intensity normalized to genetic background control. Bars represent mean ± SEM. Statistical comparisons performed using Dunn’s test with Bonferroni correction (pMad) or Tukey HSD test (TwitGFP). *** p<0.001, ** p<0.01, * p<0.05. (D) Representative images of wing phenotypes for indicated genotypes. Scale bar represents 1 mm.

**Figure 6:**
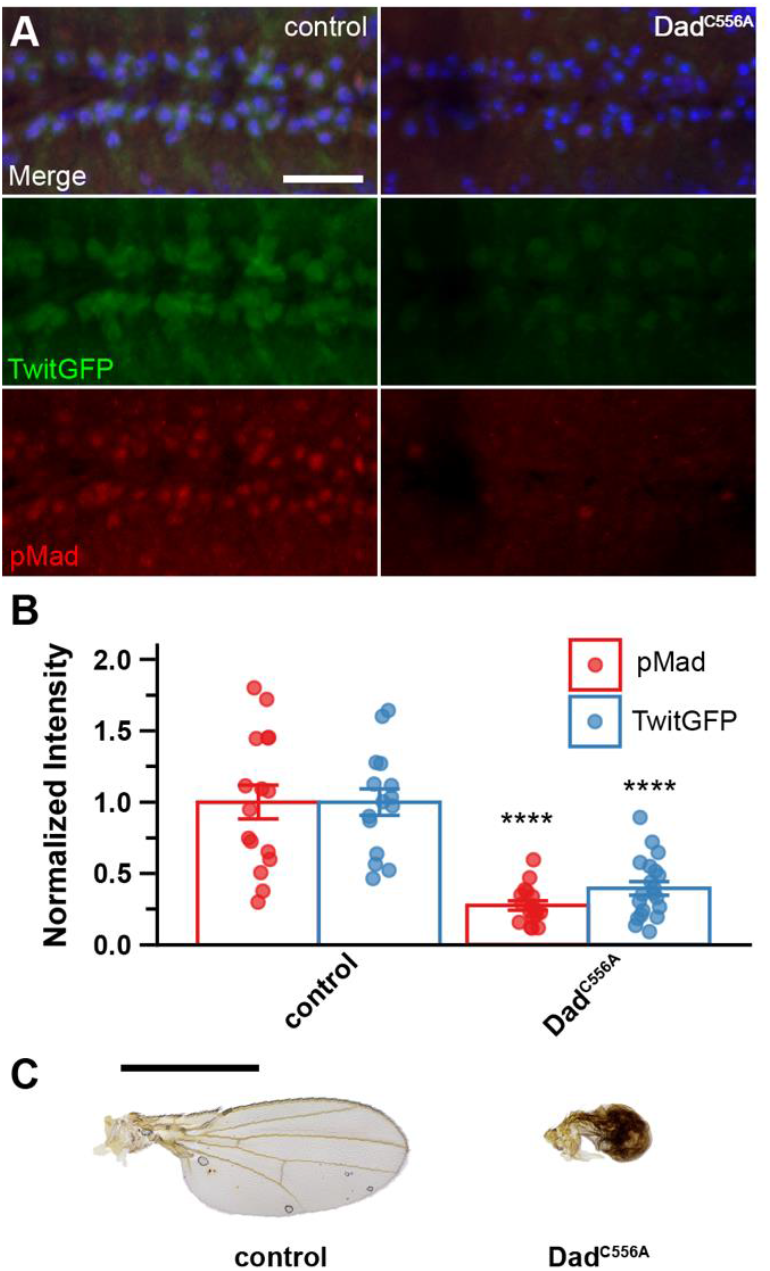
Dad functions normally without cysteine 556 in wing and brain. Transgenic control (*TwitGFP, OK371-Gal4/+*) compared with *OK371-Gal4*-driven overexpression of *UAS-Dad*^*C556A*^. (A) Motor neurons in ventral nerve cord triple-labeled for Elav (blue), Twit-GFP (green) and pMad (red). Scale bar represents 20 microns. (B) Quantification of pMad and Twit-GFP intensity normalized to genetic background control. Bars represent mean ± SEM. Statistical comparisons performed using Dunn’s test with Bonferroni correction. **** p<0.001. (C) Representative images of wing phenotypes for indicated genotypes. Scale bar represents 1 mm.

### Removal of C-terminal palmitoylation site

Previous studies identified a conserved C-terminal cysteine (C556) of Dad that serves as a palmitoylation site required for membrane association and BMP signaling inhibition (Li et al. 2017). We generated a *Dad* construct containing a substitution of C556 with alanine (Dad^C556A^). This variant was expressed using *Ap-Gal4* in the wing and *OK371-Gal4* in the brain to assess tissue-specific effects on BMP signaling. In wing tissue, expression of Dad^C556A^ produced a strong phenotype that placed it into the most severe category along with Dad (Figure 1B). This finding suggests that C556 is not essential for I-Smad function in wing tissue, contrary to its function based on ovarian studies, where this same modification reduced effectiveness to levels similar to genetic background control (Li et al. 2017). Expression of Dad^C556A^ in motor neurons resulted in a dramatic reduction in both pMad and TwitGFP levels (0.275±0.0319 and 0.395±0.0469 relative to control, respectively). These results indicate that C556-dependent palmitoylation is not universally required for Dad function, and that this process may be tissue specific, rather than universally applicable to I-Smad functionality.

### Structural predictions of physical interactions between Dad and target molecules

Our results support a model where I-Smads inhibit TGF-β /BMP signaling through several distinct mechanisms, and different tissues preferentially employ these different mechanisms. As we used structural prediction to compare a putative DNA binding motifs of R-Smads and I-Smads (Figures 2A and 5A), we wanted to test whether structural predictions can also visualize the physical interactions between I-Smads and the other target proteins, R-Smads and Type-I receptors. Indeed, AlphaFold3 models support a strong, complementary conformation between Dad and the R-Smad Mad and between Dad and the Type-I receptor Thickveins (Tkv) (Figure 7). The I-Smad/R-Smad conformation with the closest fit was between two I-Smads and one R-Smad (Figure 7A). There were five C-terminal residues of Dad that formed hydrogen bonds with the MH2 domain of Mad: M520, H523, G524, M528, and R568. Structural predictions most strongly support an association between a single I-Smad and Type-I receptor (Figure 7B). We identified nine C-terminal residues of Dad that formed hydrogen bonds with the cytoplasmic region of Tkv: T406, W406, D487, L495, M528, P530, T544, I553, E561. In agreement with prior knowledge, these residues are all found within the C-terminal MH2 domain of the I-Smad.

**Figure 7:**
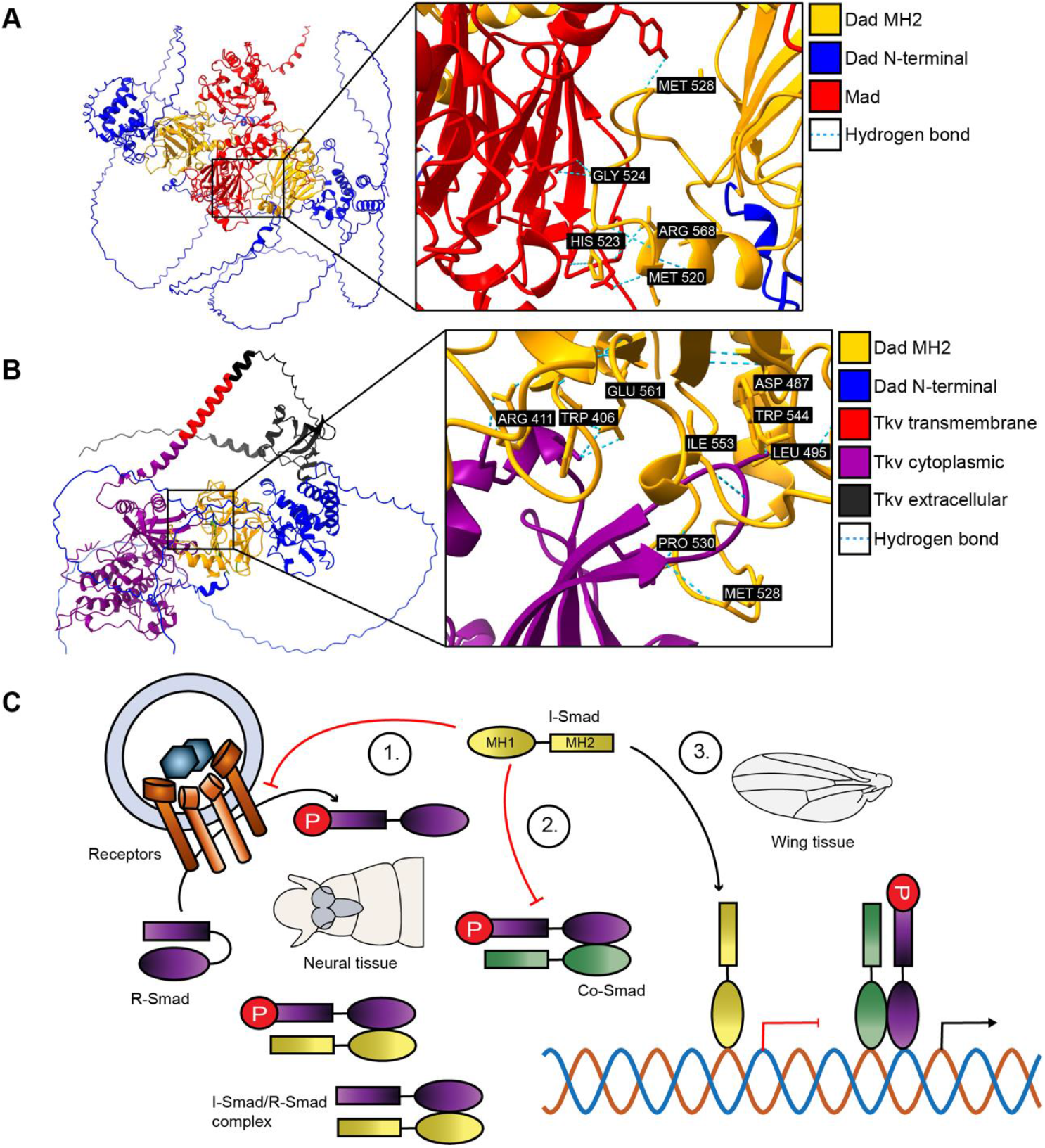
Structural predictions support multiple modes of I-Smad function. (A) AlphaFold3 predicted protein interaction between two Dad molecules (I-Smad) and one Mad (R-Smad). Predicted hydrogen bonding residues are labeled. (B) AlphaFold3 predicted protein interaction between one Dad (I-Smad) and one Thickveins (Type I receptor). Predicted hydrogen bonding residues are labeled. (C) I-Smads function through at least three distinct mechanisms: 1. Preventing receptors from phosphorylating R-Smads, 2. Binding to R-Smads and preventing formation of R-Smad/Co-Smad complex, and 3. Interacting with DNA to counteract TGF-β/BMP transcriptional effects. In neural tissue, I-Smads function preferentially through either the first or second mechanisms. In wing tissue, I-Smads rely on the third mechanism.

## Discussion

### Structural basis of I-Smad tissue-specific mechanisms

The modes by which I-Smads have been shown to inhibit TGF-β/BMP signaling can be classified into three distinct mechanisms (Figure 7C): 1. Interfering with Type I receptors, either by direct physical interaction, or by recruitment of effector molecules such as BAMBI, E3 ubiquitin ligases (Smurf1 and Smurf2), or GADD34; 2. Preventing R-Smad/Co-Smad complex formation both by direct physical interaction and promoting ubiquitination; 3. Directly regulating transcription to counteract expression changes of TGF-β/BMP target genes (Miyazawa and Miyazono 2017). Our current study supports the hypothesis that certain tissue types utilize specific I-Smad mechanisms preferentially over others. Confirming previous results, we found that expression of the *Drosophila* I-Smad, Dad, in wing primordium results in abnormally small, shriveled wings (Figure 1B) (Inoue et al. 1998). Seeking to determine which parts of Dad are sufficient for this phenotype, we identified a 24 aa deletion in a putative DNA binding region within the MH1 domain (ΔDNABD) (Figure 1A & 1B). Protein structure prediction reveals a complete disruption of the beta hairpin structure that resembles the R-Smad structure which closely contacts the DNA molecule (Figures 2A & 4A).

ΔDNABD expressed in motor neurons disrupted BMP signaling in this neural tissue (Figure 5), indicating that an alternative mechanism is used in this tissue compared to wing disc. Indeed, Dad with only an MH1 or an MH2 domain (ΔMH1 and ΔMH2) disrupted BMP signaling in neural tissue (Figure 4). However, Dad with only MH2 domain (ΔMH1), was not sufficient to disrupt wing growth (Figures 1B and 4C). Based on these results, we propose that Dad functions primarily through an MH1-mediated mechanism in this tissue, likely direct transcriptional regulation in wing tissue, but through multiple mechanisms in neural tissue (Figure 7C). To support this model, we analyzed predicted protein structures and observed three characteristics: 1. The DNA binding loop of Dad shares a similar structure with that of R-Smads (Figure 2A); 2. The structure of I-Smad associating with the Type-I receptor reveals binding between specific residues; and 3. The structure of I-Smads associating with R-Smads also reveals interaction sites (Figure 7A,B).

These alternative mechanisms may hold the key to maintaining appropriate levels of TGF-β/BMP signaling in different tissue types. For example, we found that Dad with a cysteine to alanine substitution at position 556 (C556A) was fully effective at disrupting both wing development and BMP signaling outputs in motor neurons (Figures 1B & 6). Previous work shows that this mutation abolishes the ability of Dad to cause loss of germ cells in ovarian tissue (Li et al. 2017). As this residue is reported to serve as a palmitoylation site that promotes association with receptors on the plasma membrane, it appears that Dad functions independently of this membrane association in wing and brain tissue. Further experiments such as genetic epistasis will be required to determine the exact mechanism(s) used in these tissue types.

### Multiple functions performed by distinct structural regions in other proteins

The fact that a single protein can utilize different structural regions to work through such disparate interactions is remarkable but is not without precedent. Trio, a large, multi-domain protein, exemplifies this versatility. The N-terminal GEF1, central GEF2, and C-terminal protein kinase domains each catalyze distinct reactions that affect different aspects of cellular physiology (Bircher and Koleske 2021). While GEF1 activates Rac1 to promote membrane ruffling and cell migration, GEF2 stimulates RhoA to induce stress fiber formation, with the protein kinase domain providing additional signaling capabilities. Another example is the extracellular protein Cerberus, which uses distinct mechanisms to inhibit three different pathways: directly binding to Wnt proteins in the extracellular space, preventing Nodal from interacting with its receptors, and antagonizing BMP signaling through direct protein interaction (Piccolo et al. 1999; Kawano and Kypta 2003; Tavares et al. 2007; Aykul et al. 2015). Thus, proteins can utilize different structural motifs to carry out distinct interactions, although further research is necessary to determine whether these different mechanisms are used preferentially depending on the cellular context.

### Switching I-Smad mechanisms based on intracellular environment

It is likely that the complement of molecules within a cell determines which mechanism an I-Smad will work through in that cell. Recent works have begun to elucidate the identities of I-Smad partner molecules. For example, Liu et al. recently identified a gene encoding part of the Integrator complex in a screen for genes required for female germline maintenance (Liu et al. 2023). In germline tissue and S2 cells, the Integrator complex is required for Dad to degrade Tkv. It is possible that certain Integrator complex components are absent from wing discs, and thus Dad functions through an alternative mechanism. Testing this hypothesis may prove challenging as there are eight genes encoding Integrator complex components (Hao et al. 2008).

Under the assumption that I-Smads evolved from an ancestral Smad gene that encoded a protein with DNA binding properties, we favor a model wherein *Smad6* and *Smad7* began differentiating after their duplication in the vertebrate lineage. Considering the higher percent sequence identity between Dad and Smad6, as well as the specificity of both molecules for the BMP branch of TGF-β signaling (Goto et al. 2007), it can be inferred that Smad6 retains more of the ancestral function shared with Dad. Therefore, Smad7 may have evolved additional mechanisms, for example an increased effectiveness of the MH2-mediated mechanisms (Figure 1B)(Hanyu et al. 2001), and a reduction of the MH1-mediated mechanisms. The structural similarity of the DNA binding loop between R-Smads, Smad6, and Dad, but not Smad7 (Figure 2A) supports this hypothesis.

### Applications and future directions

These findings indicate that any potential diagnosis or treatment involving I-Smads must consider the mechanism(s) employed by the target tissue type. We demonstrate that to inhibit growth of wing tissue, the *Drosophila* I-Smad functions through an MH1-mediated modality, most likely involving DNA binding. However, future work is necessary to test the ramifications and limits of this phenomenon. It is vital to determine which mechanisms work in additional tissue types, such as intestinal epithelium, and whether the mechanisms change in pathological states like colorectal cancer. Further work is also needed to directly test aspects of our model. Importantly, biochemical assays such as chromatin immunoprecipitation are needed to confirm association of I-Smads and variants with DNA. It will also be interesting to further explore the evolutionary relationships between Dad, Smad6, and Smad7 to learn more about their common and divergent functionality. Our versatile in vivo platform can be used to conduct additional structure/function analyses of I-Smad orthologs, such as deletion of the homologous Smad6

DNA binding region and domain swap experiments. Ultimately, extension of these findings to vertebrate systems will be necessary to test their broader applicability to modulating TGF-β/BMP signaling in diverse cellular contexts.

## Methods

### Molecular biology and transgenesis

Plasmids were created using standard molecular biology methods. All plasmid sequences were verified using whole plasmid nanopore sequencing (Eurofins Genomics; Louisville, KY, USA). Injections were performed by Rainbow Transgenics, Inc. (Camarillo, CA, USA). All constructs were inserted into the VK00033 locus on the third chromosome. Insertions were verified by sequencing. Complete lists of plasmids and primers used in this study are included below:

| Plasmid | Creation | Source | Notes |
|---|---|---|---|
| pUAST-attB | N/A | DGRC Stock 1419 ; https://dgrc.bio.indiana.edu//stock/1419; RRID:DGRC_1419 | Destination vector for transformation |
| pBSPPC | N/A | pBSPPc was a gift from Cuiqing Ma (Addgene plasmid # 116918 ; https://n2t.net/addgene:116918; RRID:Addgene_116918) (Xu et al. 2013) | Intermediate vector |
| CS2 Smad6 | N/A | CS2 Smad6 was a gift from Joan Massague (Addgene plasmid # 14960 ; https://n2t.net/addgene:14960; RRID:Addgene_14960) (Hata et al. 1998) | Source of Smad6 CDS |
| TFORF0111 | N/A | TFORF0111 was a gift from Feng Zhang (Addgene plasmid # 141518 ; https://n2t.net/addgene:141518; RRID:Addgene_141518) (Joung et al. 2023) | Source of Smad7S CDS |
| pBSPPc-Smad6 | Insert from CS2 Smad6 cloned into pBSPPc using BamHI/XbaI sites | Current study |  |
| pUAST-attB-Smad6 | Insert from pBSPPc-Smad6 cloned into pUAST-attB using XhoI/XbaI sites | Current study | Transformation vector for Smad6 |
| pBSPPc-NMC | Additional cloning sites added to pBSPPc by inserting oligonucleotide into PstI/BamHI sites | Current study | To accommodate Smad7S insert |
| pBSPPc-NMC-Smad7S | Insert from TFORF0111 cloned into pBSPPc-NMC using BmtI/SpeI sites | Current study |  |
| pUAST-attB-Smad7S | Insert from pBSPPc-NMC-Smad7S (BamHI/XbaI) cloned into pUAST-attB using XhoI/XbaI sites | Current study | Transformation vector for Smad7S |
| LD47465 | N/A | LD47465 (DGRC Stock 1333567 ; https://dgrc.bio.indiana.edu//stock/1333567 ; RRID:DGRC_1333567) | Source for Dad CDS |
| pUAST-attB-Dad | PCR product from LD47465 with XhoI/XbaI sites added cloned into pUAST-attB |  | Transformation vector for UAS-Dad |
| PCMV5-Smad7-HA | N/A | pCMV5-Smad7-HA was a gift from Jeff Wrana (Addgene plasmid # 11733 ; https://n2t.net/addgene:11733; RRID:Addgene_11733)(Hayashi et al. 1997) | Source for Smad7L CDS |
| PCMV5-Smad7 | Deleted HA tag from pCMV5-Smad7-HA with PCR | Current study |  |
| pUAST-attB-Smad7L | Insert from pCMV5-Smad7 cloned into pUAST-attB using EcoRI/XbaI sites | Current study | Transformation vector for UAS-Smad7L |
| pUAST-attB-Dad-ΔMH1 | Deleted | Current study | Transformation vector for ΔMH1 |
| pUAST-attB-Dad-ΔDNA-BD |  | Current study | Transformation vector for ΔDNABD |
| pUAST-attB-Dad-ΔMH2 |  | Current study | Transformation vector for ΔMH2 |
| pUAST-attB-Dad-ΔTrimer |  | Current study | Transformation vector for ΔTrimer |

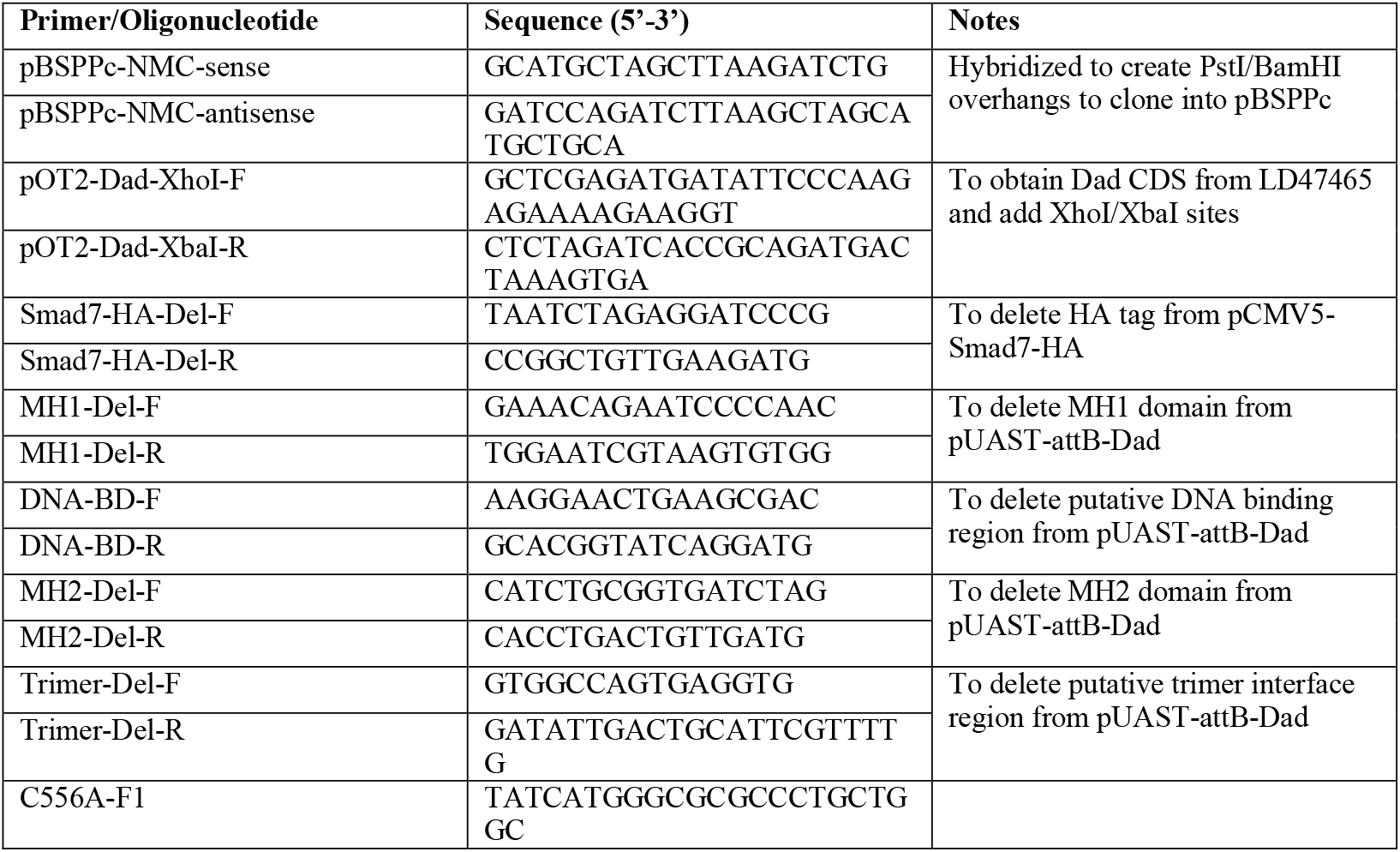
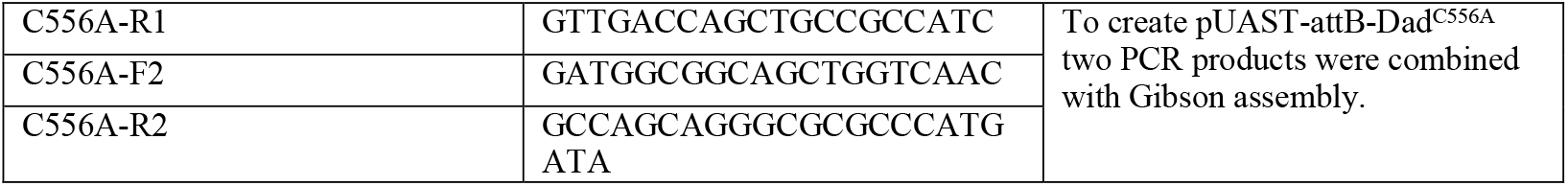

### Drosophila stocks and husbandry

Animals were reared at 25°C on standard cornmeal molasses medium. To control for crowding, crosses were set up with eight females crossed with six males, and adults were passed to a new vial every 3-4 days. For expression in motor neurons, *OK371-Gal4* (Mahr and Aberle 2006) was used. For expression in wing primordium, *Ap-Gal4* was used. A complete list of the strains used for this study is included below:

| **Abbreviation** | **Genotype** | **Source** |
|---|---|---|
| *w1118* | *w[1118]* | BIN: 2376 (BDSC) |
| *OK371* | *w[1118]; P{w[+mW.hs]=GawB}VGlut1[OK371]* | BIN 26160 (BDSC) |
| *Ap-Gal4* | *w[1118]; P{y[+t7.7] w[+mC]=GMR42A06-GAL4}attP2* | BIN 41425 (BDSC) |
| *Dad<sup>RNAi</sup>* | *y[1] v[1]; P{y[+t7.7] v[+t1.8]=TRiP.JF02133}attP2* | BIN: 26235 (BDSC) |
| *TwitGFP* | *w[1118]; Mi{y[+mDint2]=MIC}twit[MI06552]* | BIN: 41449 (BDSC) |
| *UAS-Dad* | *w[1118]; +; P{UAS-Dad}VK00037* | Current study |
| *UAS-Smad6* | *w[1118]; +; P{UAS-SMAD6}VK00037* | Current study |
| *UAS-Smad7* | *w[1118]; +; P{UAS-SMAD7.1}VK00037* | Current study |
| *UAS-Smad7S* | *w[1118]; +; P{UAS-SMAD7.3}VK00037* | Current study |
| *ΔMH1* | *w[1118]; +; P{UAS-DadΔMH1}VK00037* | Current study |
| *ΔMH2* | *w[1118]; +; P{UAS-DadΔMH2}VK00037* | Current study |
| *ΔDNABD* | *w[1118]; +; P{UAS-DadΔDNABD}VK00037* | Current study |
| *ΔTrimer* | *w[1118]; +; P{UAS-DadΔTrimer}VK00037* | Current study |
| *Dad<sup>C556A</sup>* | *w[1118]; +; P{UAS-Dad<sup>C556A</sup>}VK00037* | Current study |

### Immunostaining and microscopy

Nervous system imaging and immunostaining were performed as described (Brown et al. 2018). Briefly, wandering third instar larvae were dissected in hemolymph-like-3.1 (HL3.1) saline (Feng et al. 2004), fixed in 4% paraformaldehyde diluted with 1XPBS for 20 minutes, and washed several times in wash buffer (1XPBS with 0.3% TX-100). Primary antibodies were used at the following concentrations: rabbit anti-pSmad1/5 (Cell Signaling Technology Cat# 9516, RRID:AB_491015) at 1:500, rabbit anti-pMad PS1 (Carl-Henrik Heldin Göteborg University, Sweden Cat# pMad, RRID:AB_2617125) at 1:500, chicken anti-GFP (Abcam Cat#ab13960, RRID: AB_300798) at 1:1000, and rat anti-Elav (DSHB Cat# 7E8A10, RRID: AB_528218) at 1: 500. Secondary antibodies were used at the following concentrations: goat anti-chicken Alexa Fluor 488 (Thermo Fisher Scientific Cat# A-11039, RRID:AB_2534096) at 1:500, goat anti-rabbit Alexa Fluor 568 (ThermoFisher Scientific Cat# A-11011, RRID: AB_143157) at 1:500, goat-anti-rat Alexa Fluor 647 (Thermo Fisher Scientific Cat #A-21247, RRID: AB_141778), and goat anti-HRP Alexa Fluor 488 (Jackson ImmunoResearch Labs Cat# 123-545-021, RRID:AB_2338965). Samples were processed in parallel such that all incubation times were identical. Brains were mounted in SlowFade Diamond (ThermoFisher Cat#S36972) and filets were mounted in Prolong Diamond (ThermoFisher Cat# P36961). Images were acquired with a Zeiss LSM 880 or Nikon Ti-2 confocal microscope using a 20X or 63X objective. Images were acquired and analyzed while blinded to genotype.

### Wing dissection, imaging, and analysis

Newly eclosed (between 8-24 hrs.) adults were immediately fixed in 100% ethanol. Wings were dissected and transferred to 70% ethanol to rehydrate. Wings were then arranged on a microscope slide and covered with ProLong Diamond mounting medium (ThermoFisher Scientific Cat# P36965). A coverslip was placed on top, and the slide was allowed to cure overnight covered with a small weight to flatten the preparation. Edges were sealed with nail polish. Slides were imaged with compound microscope using 20X objective. Area was determined by manually tracing the border of the wing image using the polygon tool in ImageJ and quantifying area of the resulting polygon. Images were acquired and analyzed while blinded to genotype.

### Structural prediction and analyses

Full-length protein sequences were obtained from the National Center for Biotechnology Information (NCBI) protein database (https://www.ncbi.nlm.nih.gov/protein/) and verified for completeness and accuracy. Complete amino acid sequences were submitted to AlphaFold3 server (https://alphafoldserver.com/) using default parameters(Abramson et al. 2024). Predicted structures were downloaded from AlphaFold3 in PBD format. Structural visualization was performed using UCSF ChimeraX (Meng et al. 2023). Confidence scores provided by AlphaFold3 were used to assess the reliability of predicted structural regions, with high-confidence regions (pLDDT > 70) considered reliable for structural interpretation.

### Data analysis and statistics

Pixel intensity was measured from maximum projection confocal micrographs using a custom ImageJ script (AIABS) as previously described (Brown et al. 2019). Intensity measurements were performed while blinded to genotype. All data were subjected to outlier analysis with the Z-test method with threshold ≥ 2 STDEV from the mean. Normality of data points for each genotype was tested using a Shapiro-Wilk test. Similarity of variances was tested using Levene’s test. For data sets with multiple genotypes where measurements from each genotype were normally distributed and variances were similar, differences across genotypes were tested by one-way ANOVA. As a post-hoc test, when samples were compared only to control, Dunnett’s test was performed. When all samples were compared to each other, Tukey’s test was conducted. For data that were not normally distributed or had unequal variances, differences across genotypes were tested by the Kruskal-Wallis test. Pairwise differences were tested by post-hoc Dunn’s test with Bonferroni correction. For comparing two groups with normal distributions, Student’s t-test was used. Data were analyzed with IBM SPSS Statistics (Version 27) and R (Version 4.3.2; R Core Team 2023). Graphs were created with the *ggplot2* R package (Wickham et al. 2016). Scripts used for statistical analysis and plot creation available upon request.

## Acknowledgements

This work was supported by The National Institute of Neurological Disorders and Stroke of The National Institutes of Health under Award Number R15NS116630 to M.J.S. We thank the *Drosophila* Genomics Resource Center (NIH Grant 2P40OD010949) for plasmids. We thank the Developmental Studies Hybridoma Bank, created by the NICHD of the NIH and maintained at The University of Iowa, Department of Biology, Iowa City, IA 52242, for antibodies. Molecular graphics and analyses were performed with UCSF ChimeraX, developed by the Resource for Biocomputing, Visualization, and Informatics at the University of California, San Francisco, with support from National Institutes of Health R01-GM129325 and the Office of Cyber Infrastructure and Computational Biology, National Institute of Allergy and Infectious Diseases. We thank Stuart Newfeld and Gregory Reeves for sharing *Drosophila* stocks. Thanks to Daniel Cox, Sean Grace, Nicholas Edgington, and Rachel Jeffrey for feedback and discussions. Thanks to members of the Sulkowski lab for proofreading and editing the manuscript.

## Author contributions

MJS and AMS conceived of the study. MJS, SJS, AMS, JNH, and NJG created transformation vectors. AMS, SJS and NJG conducted experiments and analyzed data. NJG conducted structural analysis. JNH isolated and sequenced genomic DNA from transformants. All authors contributed to writing the manuscript.

